# Contemporary environmental conditions decouple ecological similarity from evolutionary relatedness in vector mosquito communities

**DOI:** 10.64898/2026.09.09.750309

**Authors:** Chao Yang, Yoshihide Maekawa, Shinji Kasai, Yukiko Higa

**Affiliations:** Department of Medical Entomology, National Institute of Infectious Diseases, Japan Institute of Health Security, Toyama 1-23-1, Shinjuku, Tokyo, Japan 162-0052

**Keywords:** vector mosquitoes, distribution modelling, niche comparison, phylogenetic generalized linear mixed model, community ecology, phylogeny

## Abstract

Understanding the processes governing community assembly remains a central challenge in ecology. Although phylogenetic niche conservatism predicts that closely related species should occupy similar ecological niches, contemporary environmental conditions may weaken this relationship by promoting ecological convergence among distantly related species. Whether such decoupling occurs in vector mosquito communities remains largely unknown despite its importance for predicting disease transmission under global environmental change. Here, we integrated species distribution models, niche comparison tests, and phylogenetic generalized linear mixed models as a unified analytical framework to investigate the relative contributions of environmental filtering, species-specific environmental responses, and phylogenetic relatedness to the assembly of Japanese vector mosquito communities. Species distributions were primarily determined by climatic and land-cover variables. Although several mosquito species exhibited highly similar ecological niches, phylogenetic relatedness explained only a small proportion of variation in species occurrence. Instead, environmental filtering together with species-specific environmental responses consistently dominated community assembly across spatial scales. Our findings demonstrate that ecological similarity can become decoupled from evolutionary relatedness under contemporary environmental conditions. This study provides a community-level framework that links species distributions, niche relationships, and evolutionary history to improve vector surveillance, anticipate community reorganization under environmental change, and support evidence-based strategies for preventing mosquito-borne disease.

## Introduction

Vector-borne diseases are threatening the health of societies globally. It is reported that vector-borne diseases have caused over 700,000 deaths annually and put 80% of the world’s population at risk of one or more of them, accounting for 17% of the global burden of communicable diseases^1^. Of these, a large proportion of disease transmissions suggest that mosquitoes are to blame^2^.

Specific ecophysiological traits feature mosquitoes in the role of an ideal vector. Mosquitoes possess functional traits, such as diapause and desiccation resistance, enabling them to readily establish in diverse environments^3^. Mosquitoes feed on various blood sources and can transmit pathogens^4,5^. The overlap of habitats, host species, and phenology among mosquito species complicates the transmission cycles of disease-causing pathogens through food chains by expanding the range of pathogen reservoirs and increasing the likelihood of pathogen transfer between mosquito species, thereby heightening the unpredictability of outbreaks of mosquito-borne diseases (e.g., accelerating pathogen circulation and prolonging the transmission season)^4,6^. Given these situations, mosquitoes are essential to maintaining the life cycles of pathogens and causing human illness. For instance, West Nile virus (WNV) is sustained in an avian sylvatic cycle by maintenance mosquito species that are ornithophagic and competent for virus transmission, and then WNV is spread to humans by bridge mosquito species that are ornithophagic, anthropophagic, and competent for virus transmission^7^. Maintenance mosquitoes and bridge vectors contribute to the risk of human WNV cases in their coinciding space and time.

Mosquito-borne disease prevalence is further affected by the spatial distribution of suitable vector species. Rapid human land-use changes, such as deforestation, agriculturalization, and urbanization, and climate change may alter mosquito spread patterns by creating novel breeding habitats and food resources, increasing the risk of zoonotic pathogen spillovers^8^. Therefore, these modifications alter the vector-host relationship when vectors are introduced into a new habitat or exposed to a new host, further accelerating the geographical expansion, transmission cycle dynamics, and even the genetic evolution of pathogens, significantly increasing the intensity and incidence of mosquito-borne disease outbreaks^9,10^. Infection of the Yellow Fever virus (YFV) in South America is one such example. In their natural environment, the YFV is mainly transmitted by *Hemagogus, Sabethes* and *Aedes* mosquitoes to monkeys in the rainforest canopy. After logging and land clearing, mosquitoes followed the canopy edge to the ground, where they fed and infected humans^3^.

Mosquito distributions and interactions within/among communities determine the prevalence of pathogens, the frequency of disease outbreaks, and the severity of public health consequences. Eliminating vectors is an effective approach to protecting against mosquito-borne diseases, as no effective therapeutic drugs or preventive vaccines currently exist for most diseases^1^. Even if primary vectors are partially controlled, secondary vectors may continue to transmit, rendering it challenging to halt the outbreak entirely^6^. The complexity increases when multiple diseases are transmitted through various vectors. Understanding the ecological niches of these mosquitoes and mapping their spatial distributions are therefore essential for efficient vector control and disease risk estimation. Functional traits unequivocally influence where species are found and how communities form by shaping interactions with the environment, other species, and their range^11,12^. Therefore, investigating trait-environment associations reveals mechanisms underlying the distributions and co-occurrence patterns of the mosquito community^11^. Meanwhile, functional traits, which are not always measurable, reflect species’ evolutionary history and biogeographic processes and are typically inherited through phylogeny^13,14^. Considering the phylogenetic signals that govern these traits is imperative^15–17^. Through these approaches, we expect to unravel how assembly processes, such as environmental filtering, influence species’ distributions and co-occurrence patterns; this remains a central challenge in ecology^18,19^. These investigations further enhance our understanding of vector-pathogen relationships.

Despite substantial advances in mosquito ecology, three important knowledge gaps remain. First, species distribution studies primarily focus on individual mosquito species and therefore provide little insight into community-level assembly. Second, although niche overlaps among mosquito vectors may strongly influence pathogen circulation, whether similar ecological niches arise from shared evolutionary history or independent adaptation to comparable environments remains unresolved. Third, few studies have simultaneously evaluated species distributions, ecological niches, and phylogenetic community structure within a unified analytical framework.

To address these questions, we used Japanese vector mosquito communities as a model system and integrated species distribution models (SDMs), niche comparison tests, and phylogenetic generalized linear mixed models (PGLMMs). Specifically, we tested three hypotheses: (H1) environmental variables determined mosquito distributions through environmental filtering; (H2) mosquito species occupying similar environmental conditions exhibited significant niche overlap; (H3) if phylogenetic niche conservatism dominated community assembly, niche similarity should correspond to a strong phylogenetic signal, whereas weak phylogenetic effects would indicate that contemporary environmental filtering overrides evolutionary constraints. By explicitly evaluating these hypotheses, our study linked species distributions, niche evolution, and community assembly to provide a mechanistic framework for understanding vector communities in the face of rapid environmental change.

## Materials and Methods

### Study region

Japan is situated at the northeast tip of Asia, characterized by a narrow strip of land with mountainous and hilly areas in the central region of the islands and plains along the coast^20^. This geographical condition confers varied climates and landscapes from north to south, as well as differences between the Pacific Ocean side and the east side of the central mountain ranges, and the Sea of Japan side and the west. The general weather is mild and humid. Summer is very hot and humid across all areas except Northern Japan, while winter is moderately cold in West and South Japan and along the Pacific Ocean side, with heavy snow in Hokkaido and on the Sea of Japan side^20^. Mountainous areas covered with forests dominate the landscape, accounting for nearly 70% of the land utilization, followed by cultivated fields at 13.9% and residential areas at 4.4%^20^. The varied environments offer an ideal setting for studying the distribution of mosquitoes and their interactions within the community.

### Mosquito sampling and environmental covariates

To identify drivers of vector mosquito distribution and community assembly, occurrence data were collected from a 10-year survey conducted in Japan from 2013 to 2022. The occurrence sites were distributed across various terrains (e.g., mountainous areas, valleys, plains, and artificially transformed lands) and landscapes (e.g., forests, grasslands, cultivated areas and urban areas) from north to south. The types of habitats included were also varied (e.g., springs, creeks, vegetation near ponds, paddies, riverbeds, parks, greenlands, reefs, wetlands, temples and shrines)^21^. Adult mosquitoes were collected by a CDC-like trap with dry ice and a sweeping net, while larvae and pupae were collected by a dipper^21^. All samples were morphologically identified at the adult stage, following Tanaka et al. (1979)^22^. A total of 16 vector species were confirmed, and 367 occurrence sites were obtained after deduplicating in a range of around 1 km × 1 km (Supplementary Table 1). This step is to avoid spatial autocorrelation issues during modelling. Duplicated sites with multiple species were merged into a single site containing occurrence information for all species. For species distribution model and niche comparison analyses, data were also obtained from the literature for some species with rare occurrences (Supplementary Table 1). For phylogenetic community analysis, we further examined the relationship at 10 km × 10 km resolution, yielding 193 occurrences.

Environmental conditions required for mosquito survival and reproduction included abiotic variables related to temperature, precipitation, and land use, as well as biotic variables related to hosts (Supplementary Table 2)^23,24^. We extracted bioclimate and elevation variables from the WorldClim, with a resolution of 30 arcsec and 5 arcmin (approximately 1 km × 1 km and 10 km × 10 km), host variables from Livestock Geo-Wiki and Socioeconomic Data and Applications Center (SEDAC) with a resolution of 30 arcsec^25–28^. Host variables were then reclassified to a 5-arcmin resolution. The landcover variable was obtained from the JAXA (https://www.eorc.jaxa.jp/ALOS/jp/dataset/lulc_j.htm) at a resolution of 15 arcsec and then reclassified to the respective resolutions. The relative humidity variable was determined as described in Yang et al. (2024)^23^. All variables were clipped to the study region. To avoid collinearity issues in the models, variables were selected using variable importance analysis and the Pearson correlation coefficient. Those with Pearson correlation coefficients > 0.8 and not ecologically relevant to mosquitoes were removed^29,30^. For phylogenetic community analysis, VIFs were also assessed to select robust variables for model explanation (Supplementary Table 2).

### Species distribution and model estimates

Each species’ occurrence data was extracted, partitioned, and modelled using the ‘Kuenm’ package in R 4.5.2 within RStudio IDE (Supplementary Table 1)^31–33^. First, we built as many models as possible by combining different regularization multipliers and feature classes of the Maxent algorithm, and selected the fittest models using partial ROC, omission rate and AICc in order of priority; finally, we created the model (for details, see ref^31^). Model performance was thus estimated using the area under the receiver operating characteristic curves (AUC), partial ROC, omission rate, and AICc. Variable importance was represented by permutation importance and the jackknife test. The relationships between mosquito occurrence probability and variable ranges were illustrated in a partial plot^34^.

### Niche comparison

The distribution maps would give a blurred sense of overlap and interactions among mosquito communities, but they do not clarify whether a relationship exists. Testing niche overlap among species is crucial, as it significantly influences their interactions. Here, we used an environmental space-based niche comparison implemented in the R package ‘Humboldt’ to disentangle these concerns^35^.

To quantify the niche similarity, two tests, namely the Niche Overlap Test (NOT) and the Niche Divergence Test (NDT), were conducted. NOT asks whether the two species’ occupied niches are equivalent in their total accessible environmental space using an Equivalence statistic, while NDT asks whether the species’ occupied niches are similar in their shared accessible environmental space. Here, we set the accessible environmental distance to 5 km for all mosquitoes for computational simplicity^36^. Then, a Background statistic was applied to each test to confirm the power of the Equivalence statistic by examining if the distributions of the two species were more or less different than would be expected given the environmental differences present in the conditions available to them. For details, the Equivalence test compares the observed niche similarity of two species with that of two resampled groups from the pooled original localities. Each group contains the same number of the original species and is reshuffled several hundred times. Significance is determined by the frequency with which the observed overlap differed from that of the reshuffled overlaps. The background test compares the observed niche similarity between two species with the overlap between one species and the spatial distribution of the other after random shifts, which is also repeated several hundred times to determine significance. It measures how that shift in geography alters the occupied environmental space while preserving most of the spatial structure of the input localities, thereby retaining the nuances associated with each dataset’s spatial autocorrelation. By combining the significant states of the two tests, we can assess whether the species’ niches differed due to divergent evolution^35^.

To conduct these statistical tests, the occurrence localities and geographic environmental variables used in the species distribution models were sampled (defined by circles) and converted into environmental space using principal component analysis. Then, continuous environmental space surfaces were created using kernel density functions, and the occupied environmental space for the focal species and its environment was estimated from the resulting principal components^35^. For details of niche comparison, refer to the literature^35,37–39^.

### Phylogenetic community structure

Once we identified the overlapping states between species pairs, we aimed to find the drivers of community assembly. We should explore how environmental gradients and phylogenetic relationships influence co-occurrence patterns. Understanding mosquito species interactions with their environments could guide targeted vector control measures.

Here, we planned to test several hypotheses affecting mosquito communities using a phylogenetic generalized linear mixed model (PLGMM) implemented in the “phyr” package^40^. First, we tested whether different environments determine the specificity of mosquito species’ occurrence; then, we tested whether each species shows a specific response to environments; finally, we tested whether phylogenetic structure affects co-occurrence patterns^12^. To achieve this, we fit 5 models. M0, as a baseline, estimated the variance explained solely by differences among sites and species, without accounting for environmental effects (i.e., treating species identity and sites as random effects and the phylogenetic relationship among species as the random-effects covariance matrix). M1 added environmental variables to M0 as fixed effects to test for environmental filtering. M2 included both environmental variables and species-specific environmental responses to test whether each species responds differently to the environment. M3 and M4 were fitted to test the roles of phylogenetic attraction and repulsion in the co-occurrence pattern of the mosquito community (for details, see the R script). Each model was tested on both a broad scale (i.e., at 10 km × 10 km resolution; test the assembly processes in niche comparison range) and a fine scale (i.e., at 1 km × 1 km resolution; test the assembly processes in SDM range), as assembly processes can vary with spatial scale^41–43^.

To fit these models, we first log-transformed variables with right-skewed data to improve the distribution of residuals. Then, we selected important variables used in the SDM using VIF < 10. For variables that showed a unimodal relationship with species occurrence upon graphical inspection, we included a quadratic term. Prior to analysis, we centred and scaled all quantitative predictors to unit variance (i.e., z-transformation) for simplicity of computation and interpretation. To evaluate quadratic terms in unimodal species-environment relationships, we first centred the predictor by subtracting its mean, then squared the centred values, and finally applied a z-transformation. This method improves model fitting by eliminating monotonic correlations between the linear and quadratic predictors for the same environmental variable^44^. The phylogenetic variance-covariance matrix was constructed from a phylogenetic tree inferred by IQ-TREE using CO data from Maekawa et al. (2016; Supplementary Fig. 1)^45–48^. For the results of PGLMMs, Model comparisons were estimated using the widely applicable information criterion (WAIC), and random and fixed effects were evaluated using 95% credible intervals. When the di erence in WAIC (ΔWAIC) was >5, we concluded that the model had substantially improved performance^49^.

## Results

### Distributions of vector mosquitoes

Selected models for the distributions of vector mosquitoes performed well, with AUC ranging from 0.834 to 0.998, omission rate ranging from 0 to 0.091, and W_AICc ranging from 0.303 to 1. Partial ROCs were all significant (Supplementary Table 3). These consistently high values confirm the models’ reliability and accuracy. The major factors responsible for habitat suitability and range varied by species. The most important covariates were elevation, mean temperature of the warmest quarter (bio10), minimum temperature of the coldest month (bio6), mean diurnal range (bio2), annual precipitation (bio12), and land cover. Host covariates were not ranked as a priority (Fig. 1; Supplementary Table 4 & Fig. 2). The occurrence probability of each mosquito being associated with each covariate was listed in Supplementary Fig. 3 and was also species-specific.

**Fig. 1.**
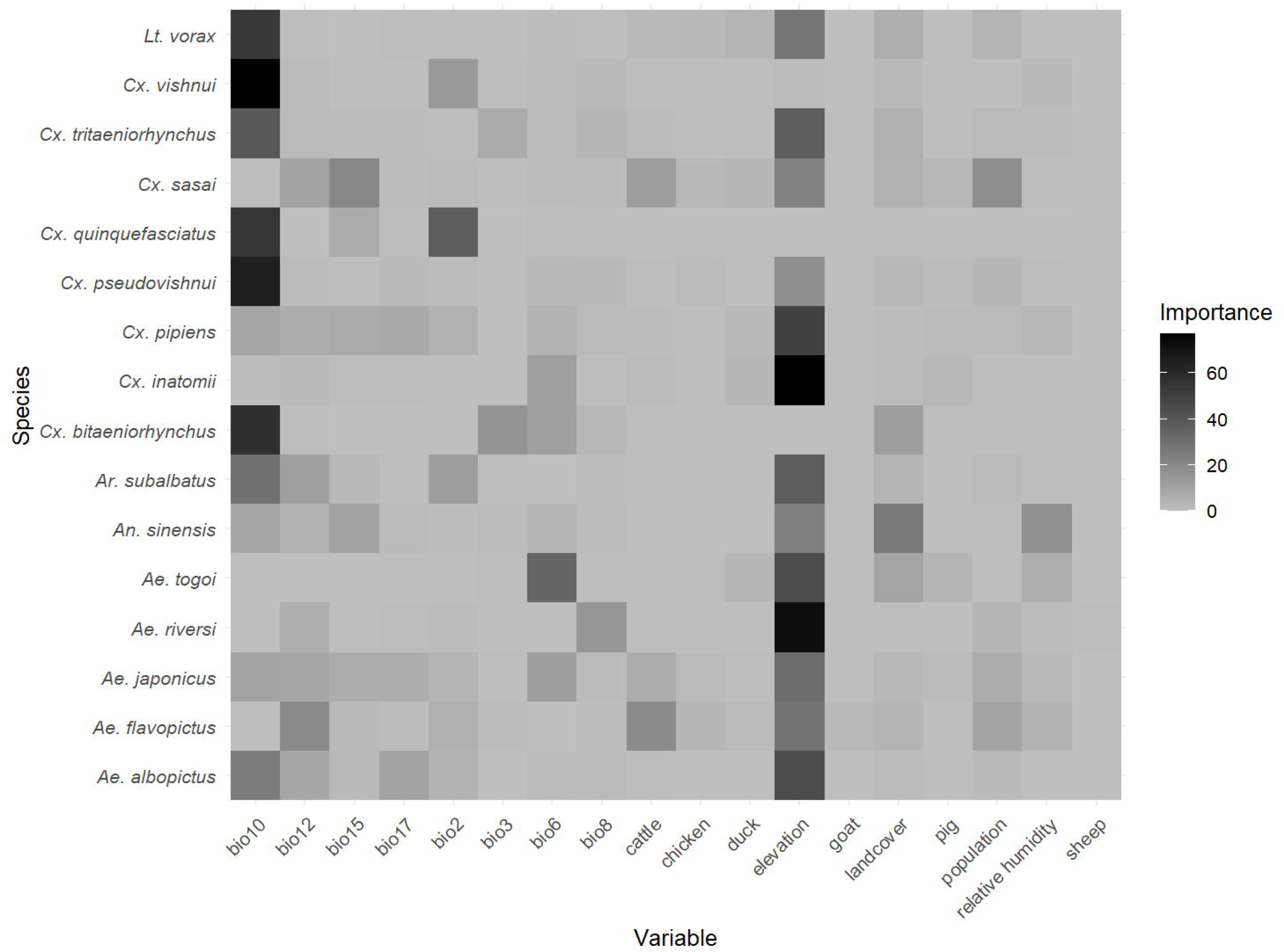
Heatmap of the variable importance for each mosquito species. For the full names of covariates, refer to Supplementary Table 2.

**Fig. 2.**
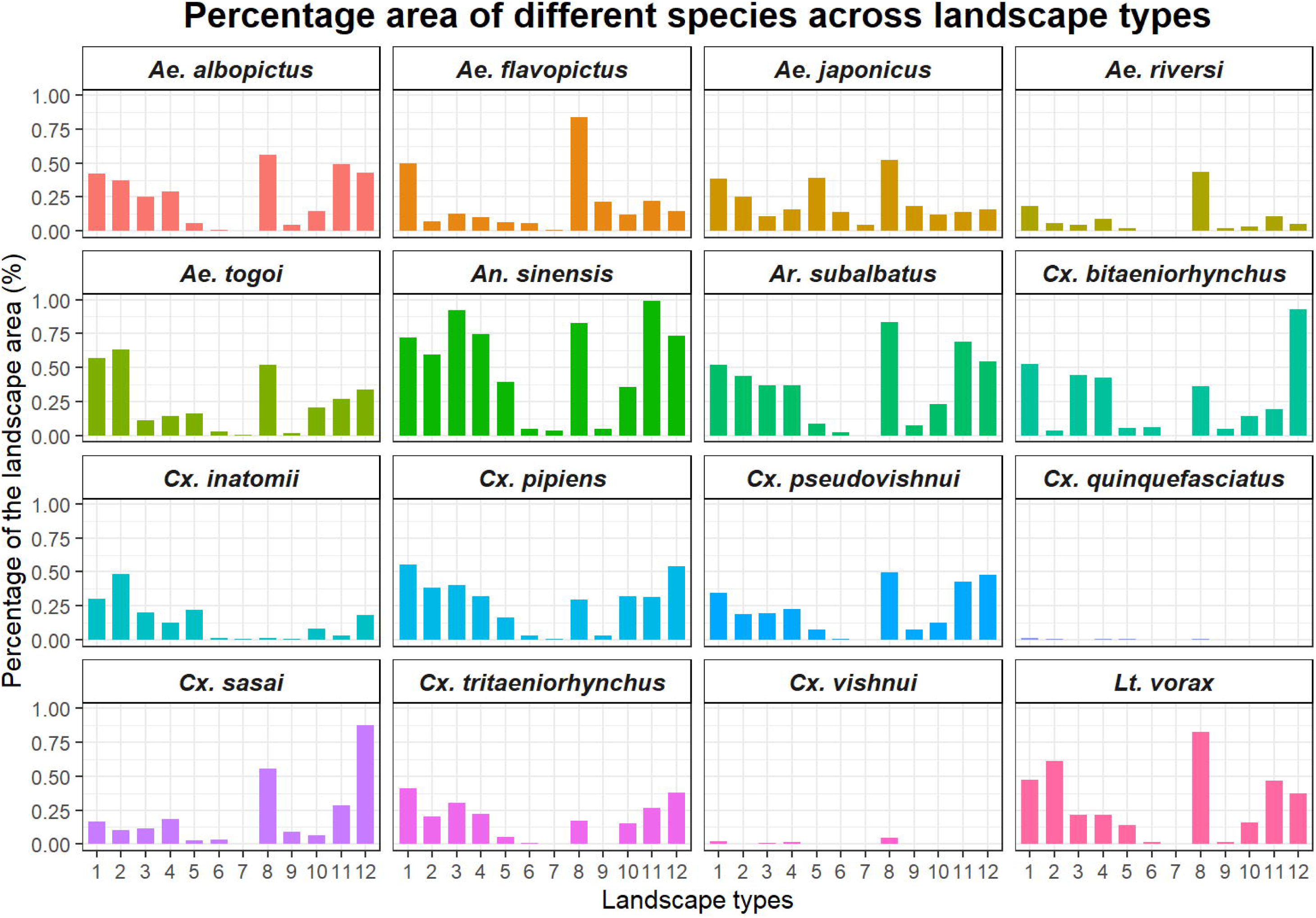
Percentage of area occupied by each mosquito species across different landscape types by establishment status. Landcover types 1-12 are water, urban, rice paddy, crop, grass, Deciduous Broadleaf Forest (DBF), Deciduous Needleleaf Forest (DNF), Evergreen Broadleaf Forest (EBF), Evergreen Needleleaf Forest (ENF), bare land, bamboo, and solar panel.

The distributions of mosquitoes’ spatial patterns were species-specific, could be defined as three types: those mainly distributed in the Ryukyu Archipelago, like *Culex vishnui* and *Culex quinquefasciatus*; those mainly distributed in Honshu Island, like *Aedes albopictus*, *Aedes riversi*, *Armigeres subalbatus*, *Culex pseudovishnui*, *Culex sasai, Culex tritaeniorhynchus*, and *Lutzia vorax*; and those distributed across the Japan Archipelago, *Aedes flavopictus*, *Aedes japonicus*, *Aedes togoi*, *Anopheles sinensis*, *Culex bitaeniorhynchus*, *Culex inatomii*, and *Culex pipiens* (Supplementary Figs. 4-5). The main land cover types utilized by mosquitoes were evergreen broad-leaved forests, bamboo groves, urban areas, paddy fields, croplands, and solar panels. *Ae. albopictus, Ae. togoi, Cx. inatomii, Cx. quinquefasciatus*, and *Lt. vorax* were more frequently appearing in the urban land use. On the other hand, *An. sinensis*, *Cx. bitaeniorhynchus*, and *Cx tritaeniorhynchus* were more often living in the paddy field. *Cx. pipiens*, *Cx. pseudovishnui*, and *Ar. Subalbatus* were frequently found in both areas (Fig. 2).

### Niche overlapping of vector mosquitoes

Pairwise comparisons of mosquito ecological niches showed that most species had either divergent niches from other species or non-equivalent niches due to differences in accessible environments, as indicated by the NOT and NDT tests (Table 1). Among these results, several mosquito pairs were suggested to occupy equivalent niches. For details, *Ae*. *albopictus* occupied a niche analogous to those of *Cx*. *pipiens* (Equivalence test: NOT [Schoener’s D = 0.738, *p* > 0.05], NDT [Schoener’s D = 0.743, *p* > 0.05]), *Cx*. *sasai* (NOT [Schoener’s D = 0.715, *p* > 0.05], NDT [Schoener’s D = 0.724, *p* > 0.05]), and *Cx*. *tritaeniorhynchus* (NOT [Schoener’s D = 0.378, *p* > 0.05], NDT [Schoener’s D = 0.676, *p* > 0.05]). It had been suggested that *Ae*. *flavopictus* possesses a similar ecological niche to *Ae*. *japonicus* (NOT [Schoener’s D = 0.773, *p* > 0.05], NDT [Schoener’s D = 0.498, *p* > 0.05]). Additionally, *Ae. japonicus* shared a comparable ecological niche with *Cx*. *sasai* (NOT [Schoener’s D = 0.819, *p* > 0.05], NDT [Schoener’s D = 0.597, *p* > 0.05]). *Ae*. *togoi* exhibited an equivalent niche to *Cx*. *pipiens* (NOT [Schoener’s D = 0.934, *p* > 0.05], NDT [Schoener’s D = 0.472, *p* > 0.05]) and *Cx*. *pseudovishnui* (NOT [Schoener’s D = 0.608, *p* > 0.05], NDT [Schoener’s D = 0.588, *p* > 0.05]). *An. sinensis* had a similar niche with *Cx. pipiens* (NOT [Schoener’s D = 0.919, *p* > 0.05], NDT [Schoener’s D = 0.495, *p* > 0.05]) and *Cx. Tritaeniorhynchus* (NOT [Schoener’s D = 0.514, *p* > 0.05], NDT [Schoener’s D = 0.623, *p* > 0.05]). *Cx*. *bitaeniorhynchus* occupied an ecological niche akin to *Cx*. *sasai* (NOT [Schoener’s D = 0.558, *p* > 0.05], NDT [Schoener’s D = 0.628, *p* > 0.05]). Furthermore, *Cx*. *pipiens* demonstrated a comparable niche to *Cx*. *pseudovishnui* (NOT [Schoener’s D = 0.668, *p* > 0.05], NDT [Schoener’s D = 0.748, *p* > 0.05]) and *Cx*. *tritaeniorhynchus* (NOT [Schoener’s D = 0.675, *p* > 0.05], NDT [Schoener’s D = 0.652, *p* > 0.05]). Lastly, *Cx*. *pseudovishnui* held an ecological niche similar to that of *Cx*. *sasai* (NOT [Schoener’s D = 0.61, *p* > 0.05], NDT [Schoener’s D = 0.786, *p* > 0.05]).

**Table 1.**
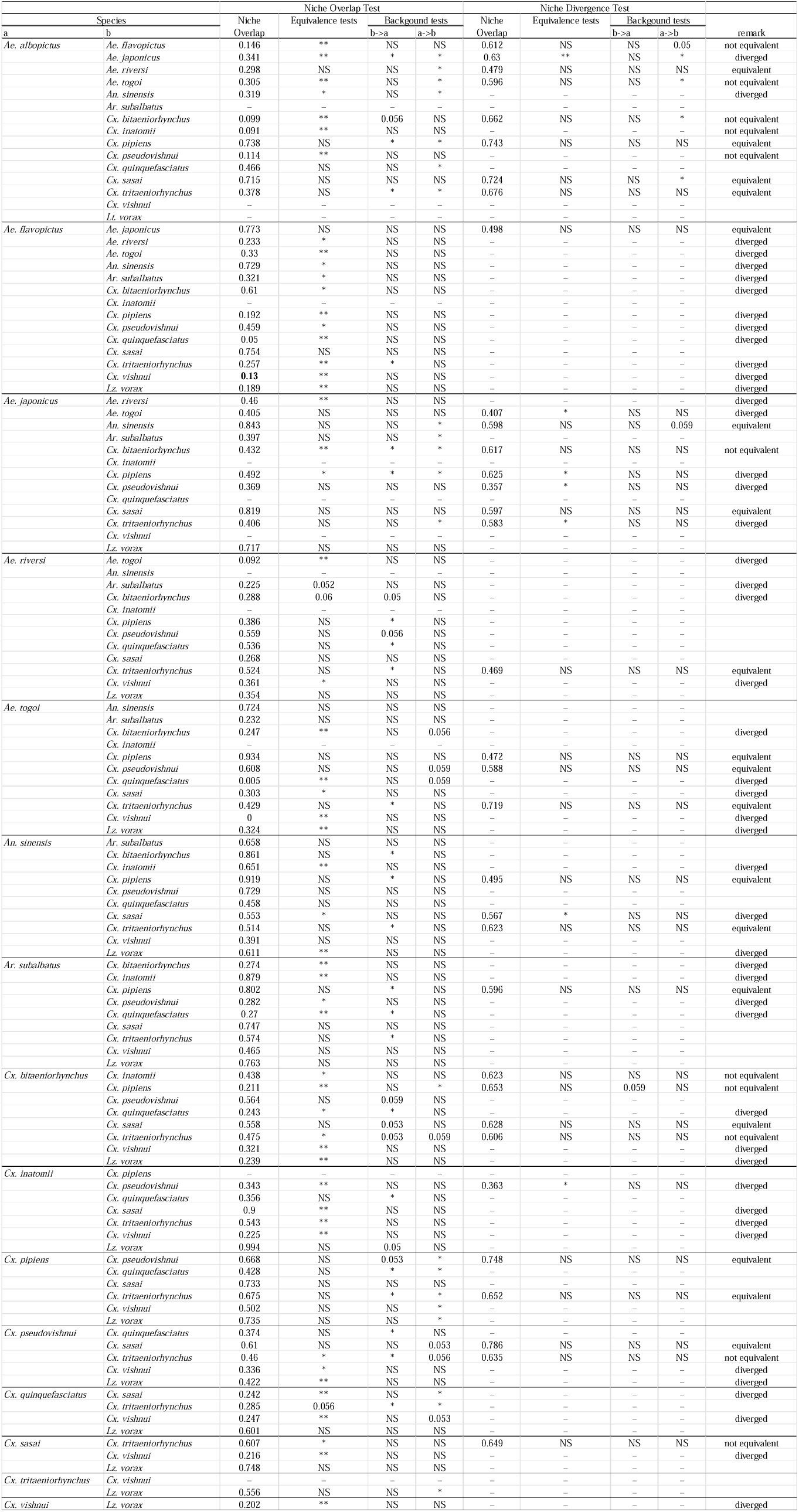
Niche overlap tests and divergence tests of mosquito pairs. Schoener’s D represents the degree of similarity between the niches of two species, ranging from 0 to 1. A value of 1 indicates niche equivalence, whereas a value of 0 indicates complete niche differentiation^61,62^. b -> a and a -> b corresponded to Background statistics comparing species b to species a or species a to simulated species b, respectively. In both cases, the first-listed species is the one whose range has shifted. Test significance: *: 0.01-0.05, **: < 0.01.

### Phylogenetic community structure of vector mosquitoes

In both scales of mosquito habitat, the best PGLMM models were M3 (WAIC = 3586 at the 1 km × 1 km scale; WAIC = 2061 at the 10 km × 10 km scale). These two models, compared to the other models on their own scales, had a significant ΔWAIC, indicating that the roles of environments, species-specific responses, and phylogenetic attraction in the distributions and co-occurrences of mosquito communities (Table 2). Here, we presented the M3 results at the 1 km × 1 km scale, as it performed well and incorporated more information (Supplementary Table 5).

**Table 2.**
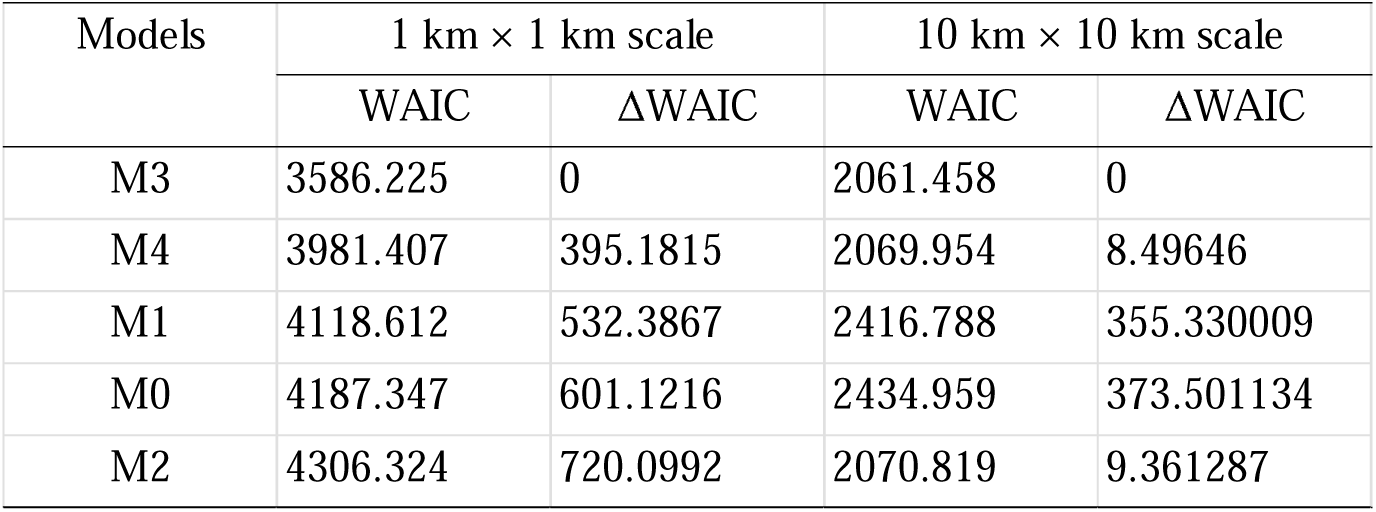
Summary information for models quantifying the strength of environmental filtering, species-specific response, and phylogenetic relationships at two spatial scales.

| Models | 1 km × 1 km scale |  | 10 km × 10 km scale |  |
| --- | --- | --- | --- | --- |
|  | WAIC | ΔWAIC | WAIC | ΔWAIC |
| M3 | 3586.225 | 0 | 2061.458 | 0 |
| M4 | 3981.407 | 395.1815 | 2069.954 | 8.49646 |
| M1 | 4118.612 | 532.3867 | 2416.788 | 355.330009 |
| M0 | 4187.347 | 601.1216 | 2434.959 | 373.501134 |
| M2 | 4306.324 | 720.0992 | 2070.819 | 9.361287 |

Random effects allow us to assess how much of the among species variation is influenced by phylogenetic relatedness (Fig. 3; Supplementary Table 6). Accordingly, these effects were formulated as non phylogenetic species effects (|sp), phylogenetically structured species effects (|sp), and the species by site interaction (sp @site). The variance of the overall nonphylogenetic species effect (1|sp) was 2.316, the largest in all random terms, whereas the phylogenetic component (1|sp) was only 0.027. The site□level variance (1|site) was tiny (0.014), whereas the species□by□site interaction (1|sp @site) variance was moderate (0.215). For among□species variation in environmental responses, sheep density showed the largest species□level slope variance (the non phylogenetic component is 2.115), while the phylogenetic component was only 0.015. Elevation and human population also had relatively high non□phylogenetic slope variances (0.164 and 0.182, respectively). In contrast, the phylogenetic slope variance for annual precipitation (bio12), cattle, chicken, goat and duck densities were comparatively higher than their non phylogenetic counterparts. For most predictors, the posterior credible intervals of phylogenetic variances were wide.

**Fig. 3.**
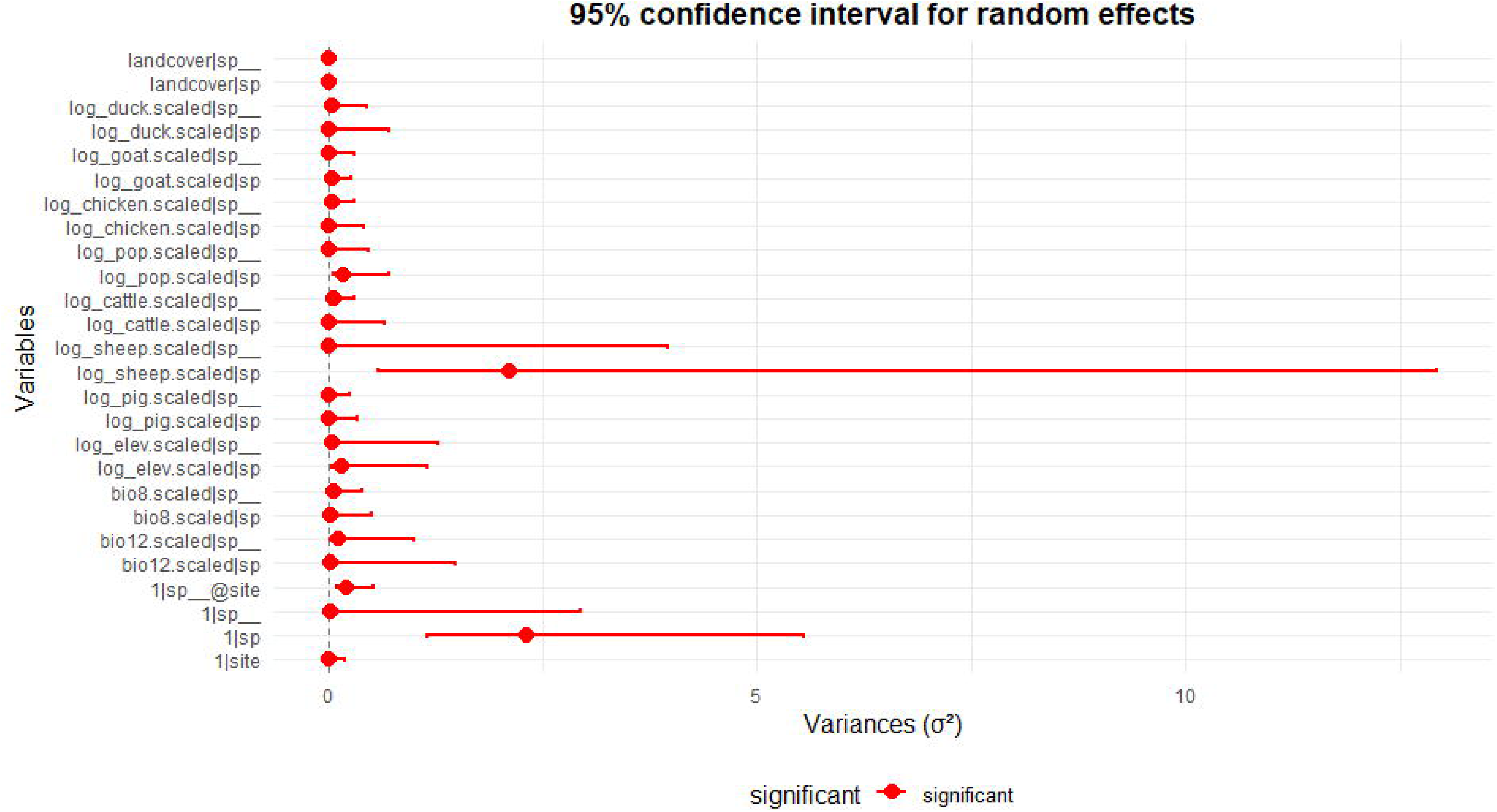
Approximate marginal posterior distribution of random effects with 95% credible interval.

The fixed effects indicate the average linear or non□linear relationships between predictors and the probability of occurrence, assessed by whether the 95% credible interval excludes zero (Fig. 4; Supplementary Table 7). Quadratic term of annual precipitation (bio12.squared. scaled; model estimate = −0.350, CI [−0.486, −0.215]), Sheep density (log_sheep.scaled; model estimate = −1.045, CI [−1.883, −0.145]), and urban land□cover (landcover2; model estimate = −0.644, CI [−1.246, −0.043]) had a significant negative effect on the occurrence probabilities of mosquito species. Quadratic term of chicken density (log_chicken.squared.scaled; model estimate = 0.180, CI [0.034, 0.327]) and solar panel land□cover (landcover12; model estimate = 1.462, CI [0.151, 2.774]) had a significant positive effect on the occurrence probabilities of mosquito species. Temperature related variables (bio8 and its quadratic term), densities of pig, cattle, goat, duck, human population (linear and quadratic), elevation, and most other land□cover types (e.g., landcover3–11) had global average effects that do not reach significance since the 95% credible interval across zero.

**Fig. 4.**
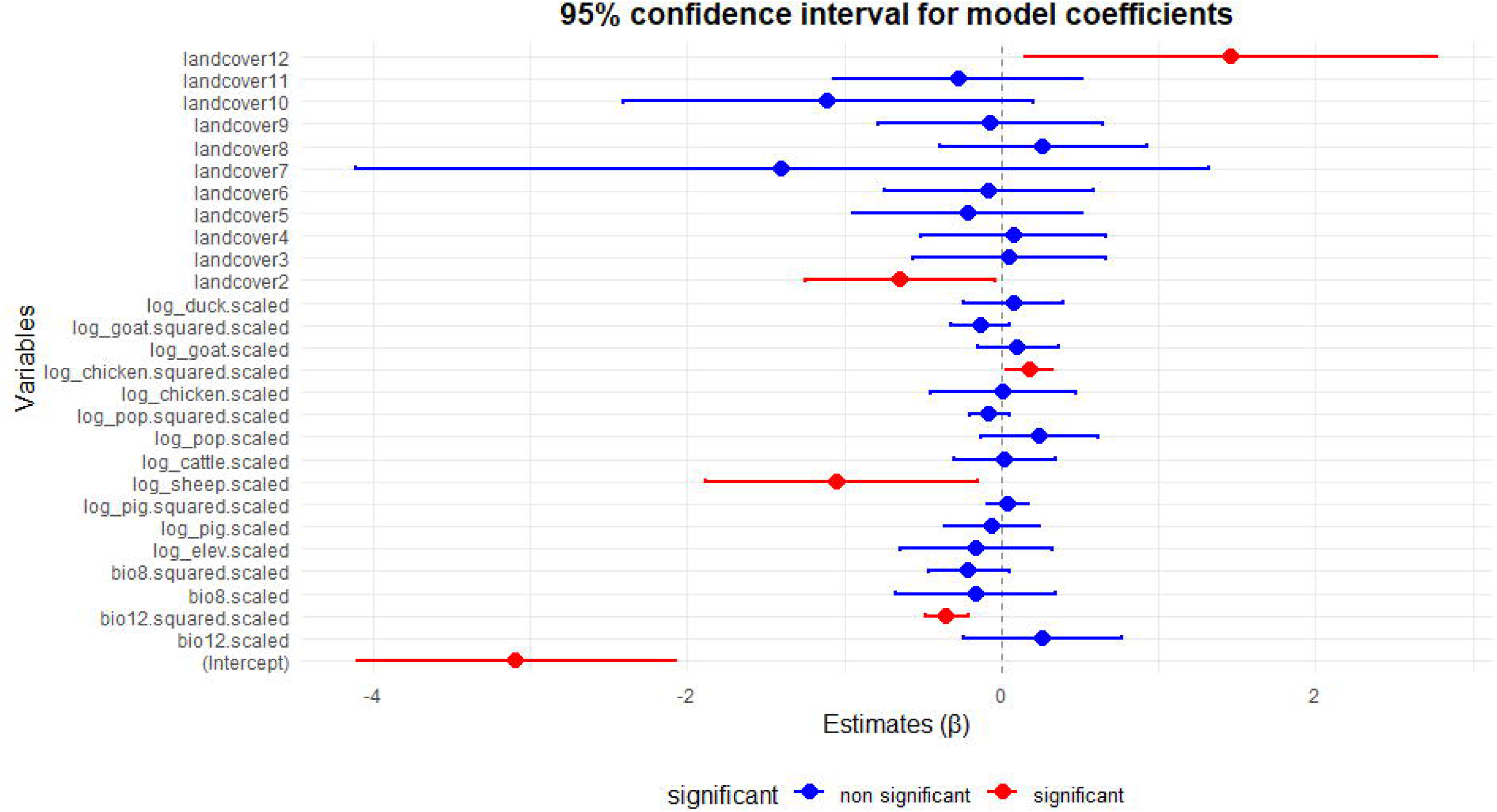
Approximate marginal posterior distribution of fixed effects with 95% credible interval. Blue: non-significant; red: significant.

## Discussion

### Environmental filtering dominates mosquito community assembly

Our results indicate that environmental filtering is the dominant process structuring vector mosquito communities in Japan. Across SDMs and PGLMMs, climatic and landscape variables showed strong associations with mosquito occurrence, whereas phylogenetic variance was comparatively weak. This supports the view that environmental conditions constrain local species establishment and persistence from a broader regional species pool^42,50^. The SDMs identified climate, elevation, and land-cover variables as important predictors of mosquito occurrence (Figs. 1-2; Supplementary Fig. 2). Temperature and precipitation directly influence mosquito development, survival, reproduction, and the availability of breeding habitat, while land cover modifies habitat structure and local environmental conditions^51,52^. Although the importance and direction of individual predictors differed among species (Supplementary Fig. 3), their overall contribution indicates that contemporary environmental conditions strongly constrain mosquito distributions. The PGLMM results further demonstrate that these environmental associations are not strongly structured by evolutionary history. The non-phylogenetic species intercept variance was 2.316, compared with only 0.027 for the phylogenetic component. Similarly, phylogenetic random-slope variances were generally small relative to species-specific variation (Fig. 3; Supplementary Table 6). Thus, after accounting for environmental variation, much of the remaining heterogeneity among species was not explained by phylogenetic relatedness. Environmental filtering therefore appears strong, but it operates primarily through species-specific ecological responses rather than conserved responses among closely related taxa^11,50^.

### Ecological similarity is decoupled from phylogenetic relatedness

Several pairs of mosquito species showed substantial environmental niche overlap, including *Ae. albopictus*–*Cx. pipiens* (Schoener’s D = 0.738), *Ae. flavopictus*–*Ae. japonicus* (D = 0.773), and *Ae. togoi*–*Cx. pipiens* (D = 0.934; Table 1). These results indicate that distantly related mosquito species can occupy similar environmental spaces. This pattern provides limited support for strong phylogenetic niche conservatism (PNC), which predicts that closely related species tend to retain similar ecological characteristics inherited from common ancestors^53^. If PNC were the dominant mechanism, ecological similarity should generally increase with phylogenetic relatedness. Instead, our PGLMMs showed relatively weak phylogenetic covariance even among species with high niche overlap (Fig. 3; Supplementary Table 6). This suggests that ecological similarity and evolutionary relatedness are partly decoupled in the mosquito community. Ecological convergence provides one possible explanation. Distantly related species may independently exploit similar contemporary environments when exposed to comparable climatic and landscape conditions^54^. Human-modified habitats, artificial water bodies, and urban or agricultural environments may create recurring ecological opportunities that can be exploited by species with different evolutionary histories. Similar environmental conditions can therefore favour similar ecological distributions among unrelated taxa, producing ecological convergence without necessarily implying close phylogenetic relatedness^54^. However, our data cannot distinguish genetic adaptation from phenotypic plasticity; therefore, convergence here refers to similarity in ecological responses rather than demonstrated convergent evolution. High niche overlap also does not imply ecological equivalence. Species occupying similar environmental space may differ in larval habitat use, host association, phenology, or responses to particular environmental gradients. Such differences can generate niche differentiation and promote coexistence despite substantial overall niche overlap^55,56^. Thus, the combination of high environmental overlap and weak phylogenetic structure may reflect ecological convergence at broad environmental dimensions together with differentiation along specific ecological axes.

### Species-specific environmental responses facilitate coexistence

The large variation in species-specific environmental responses provides an important mechanism underlying this pattern. PGLMM random slopes quantify how the strength and direction of associations between environmental predictors and occurrence vary among species (Fig. 3; Supplementary Table 6; the strength and direction of environment-occurrence associations of each species referring to Supplementary Fig. 3). Sheep density showed the largest non-phylogenetic random-slope variance (2.115), while human population and elevation also showed substantial species-level variation (0.182, 0.164). In contrast, the corresponding phylogenetic slope variances were very small, with values of 0.015, 0.007, and 0.040 for sheep, human density, and elevation, respectively. These results indicate that mosquito species respond heterogeneously to the same environmental gradients and that this heterogeneity is weakly predicted by phylogenetic relatedness. Importantly, the contrast between SDMs and PGLMMs does not reflect differences in spatial scale because both analyses used the same spatial units and environmental predictors. Rather, the two approaches address different aspects of species–environment relationships. SDMs characterize associations between environmental predictors and occurrence for individual species, whereas PGLMMs additionally quantify community-level mean effects, species-specific response variation, and phylogenetic covariance. Consequently, a relatively weak average effect can coexist with substantial interspecific variation when species respond differently to the same environmental gradient (Figs. 1-4; Supplementary Figs. 2-3 & Tables 6-7). This distinction is particularly important for host-associated variables. Although livestock and human-related predictors may have weaker average effects than climatic variables, their large species-specific slope variances suggest that host-associated environments can differentiate mosquito species. Such variation may reflect differences in host-seeking behaviour, blood-feeding patterns, or reproductive ecology^57^. However, livestock density should be interpreted as a proxy for host availability or host-associated environments, rather than as direct evidence of host preference, because host choice was not measured directly. Species-specific response heterogeneity may also facilitate coexistence. Two species can occupy similar overall environmental space while responding differently to individual environmental gradients, reducing ecological equivalence and allowing multiple taxa to persist under similar conditions^55^. Phenotypic plasticity may further contribute to such heterogeneity by allowing species to adjust their ecological performance across environmental conditions^58^. Distinguishing plasticity from genetically based adaptation will require experimental and genomic studies.

### Implications for vector ecology and disease transmission

The strong influence of climate and land use has important implications for vector ecology under environmental change. Because mosquito species respond differently to environmental gradients, climate and land-use change are unlikely to affect all vectors uniformly. Instead, environmental change may generate species turnover and novel combinations of vector species. Urbanization is particularly important because urban landscapes contain heterogeneous combinations of artificial containers, vegetation, drainage systems, and human-associated hosts. Our results indicate that human population density and urban-related land cover are associated with mosquito occurrence, but their effects are species-specific rather than universally positive (Figs. 3-4; Supplementary Tables 6-7). Urbanization should therefore not be treated simply as a proxy for increased mosquito richness. Solar panel-associated land cover provides an emerging example (Fig. 4; Supplementary Table 7). Solar panel sites showed a positive association with mosquito occurrence in the community model, suggesting that photovoltaic landscapes may provide conditions conducive to certain mosquito species. However, this association does not demonstrate that solar panel sites increase mosquito breeding or pathogen transmission. Field studies are needed to determine whether photovoltaic infrastructure provides suitable breeding habitats or correlates with other environmental characteristics. The epidemiological consequences should also be interpreted cautiously. Mosquito occurrence alone does not determine disease transmission, which depends on vector abundance, competence, host availability, pathogen prevalence, and biting behaviour. Environment-driven redistribution of mosquito species may nevertheless alter the spatial context of arbovirus transmission^52^. Community-level surveillance that integrates species identity, environmental suitability, host-associated variables, and pathogen surveillance may therefore provide a more informative assessment of future vector risk than single-species monitoring alone.

### Limitations and future directions

Several limitations should be considered when guiding future directions. First, mosquito phylogeny was primarily based on COI, which provides useful species-level information but may not fully represent genome-wide evolutionary relationships. Genome-scale or multilocus phylogenies would provide a stronger test of whether the weak phylogenetic structure observed here persists across the genome^41^. Second, the lack of detailed functional trait data limits our ability to identify the mechanisms underlying species-specific responses^11^. Integrating thermal tolerance, larval habitat preference, oviposition preference, host use, and diapause would allow direct tests of whether ecological convergence reflects shared functional strategies^59,60^. Population genomic data would further help distinguish phenotypic plasticity from genetically based adaptation. Finally, our analyses describe contemporary species–environment associations and therefore cannot directly predict future community composition. Integrating species-specific responses with CMIP6 climate projections and future land-use scenarios would allow prediction of community turnover, species richness, and taxonomic beta diversity. Such community-level projections could reveal where climate and land-use change are most likely to generate novel vector assemblages.

## Conclusions

Our results support three main conclusions. First, environmental filtering is the dominant process structuring mosquito communities. Second, ecological similarity is partly decoupled from phylogenetic relatedness, indicating limited support for strong phylogenetic niche conservatism. Third, species-specific environmental responses are substantially stronger than phylogenetically structured responses, providing a potential mechanism for ecological differentiation and coexistence. We therefore propose a species-specific environmental filtering framework for mosquito community assembly. Contemporary climate and land use first filter the regional species pool; species-specific responses then determine community composition within environmentally suitable conditions; and differentiation in responses to particular environmental gradients may facilitate coexistence among species sharing broadly similar niches. Phylogenetic relatedness imposes comparatively weak constraints on these contemporary responses. This framework suggests that predicting future vector communities requires moving beyond phylogenetic relatedness as a proxy for ecological similarity and explicitly incorporating species-specific environmental responses, functional traits, evolutionary history, and future climate and land-use change. Such an approach may improve our ability to anticipate mosquito community reorganization and its potential consequences for mosquito-borne disease transmission.

## Acknowledgment

This work was supported by the Ministry of Health, Labour and Welfare of Japan (Grant Number: H24-Shinko-Ippan-007), and the Japan Agency for Medical Research and Development (Grant Numbers: JP17fk0108311, JP20fk0108067, JP23fk0108613, JP24fk0108693 and JP26fk0108693).

## Author Contributions

C.Y. designed research; C.Y. performed research; C.Y., Y.M. collected data;

C.Y. analysed data; C.Y. wrote the paper; and C.Y., Y.M., S.K., and Y.H. reviewed the paper.

## Data Availability

Datasets and codes are accessible through the link below: https://doi.org/10.57760/sciencedb.0122l.

## Figure and Table legends

Supplementary Fig. 1 Phylogeny of the 16 mosquito species. This phylogenetic tree is constructed using the IQ-TREE software based on the Maximum Likelihood model. The optimal evolutionary model is determined using the “-m MFP” and “-mtree” commands, *An. sinensis* is designated as the outgroup, and 1,000 ultrafast bootstrap replicates are performed.

Supplementary Fig. 2 Jackknife tests of variable importance of 16 mosquito species. Dark blue bars represent the predictive contribution of each variable when used alone. Light blue bars reveal how much information is unique to a variable, showing the performance drop when that variable is omitted. The red bar represents the performance of the full model, including all predictive variables.

Supplementary Fig. 3 Partial plots of 16 mosquito species showing the relationships between environmental predictors and the predicted probability of occurrence. Red curves illustrate the mean response when a variable is changed while holding all other environmental variables at their average values. Blue shadings represent the mean standard deviation. Values on the Y-axis represent the probability of occurrence.

Supplementary Fig. 4 Distribution maps of 16 vector mosquito species in Japan. The deeper the black, the higher the probability of occurrence. (A). *Ae. albopictus*, (B). *Ae. falvopictus*, (C). *Ae. japonicus*, (D). *Ae. riversi*, (E). *Ae. togoi*, (F). *An. sinensis*, (G). *Ar. subalbatus*, (H). *Cx. bitaeniorhynchus*, (I). *Cx. inatomii*, (J). *Cx. pipiens*, (K). *Cx. pseudovishnui*, (L). *Cx. quinquefasciatus*, (M). *Cx. sasai*, (N). *Cx. tritaeniorhynchus*, (O). *Cx. vishnui*, (P). *Lt. vorax*.

Supplementary Fig. 5 Binary maps for the establishment status of 16 vector mosquito species in Japan. The binary maps were generated using logistic thresholds for maximum training sensitivity and specificity in the prediction models. They are 0.2833 for *Ae. albopictus* (A), 0.5012 for *Ae. falvopictus* (B), 0.3483 for *Ae. japonicus* (C), 0.2777 for *Ae. riversi* (D), 0.3458 for *Ae. togoi* (E), 0.3828 for *An. sinensis* (F), 0.2529 for *Ar. subalbatus* (G), 0.4943 for *Cx. bitaeniorhynchus* (H), 0.1251 for *Cx. inatomii* (I), 0.3096 for *Cx. pipiens* (J), 0.1902 for *Cx. pseudovishnui* (K), 0.2594 for *Cx. quinquefasciatus* (L), 0.4267 for *Cx. sasai* (M), 0.3146 for *Cx. tritaeniorhynchus* (N), 0.1124 for *Cx. vishnui* (O), 0.2629 for *Lt. vorax* (P).

Supplementary Table 1. Occurrence information for 16 mosquito species.

Supplementary Table 2. Covariates used in SDMs, niche comparison, and PGLMMs.

†: Covariates used for phylogenetic generalized linear mixed model.

Supplementary Table 3. Metrics used to validate the outputs of species distribution models for 16 mosquito species.

†: M for regularization multipliers; F for feature classes, l = linear, q = quadratic, p = product, t = threshold, and h = hinge.

Supplementary Table 4. Variable importance for each mosquito species

Supplementary Table 5. Metrics used to validate the output of the phylogenetic generalized linear mixed model for 16 mosquito species.

Supplementary Table 6. Approximate marginal posterior distribution of random effects with 95% credible interval.

Supplementary Table 7. Approximate marginal posterior distribution of fixed effects with 95% credible interval.

