## Supplementary figures and images for "Contemporary environmental conditions decouple ecological similarity from evolutionary relatedness in vector mosquito communities"

### Supplementary Fig. 1

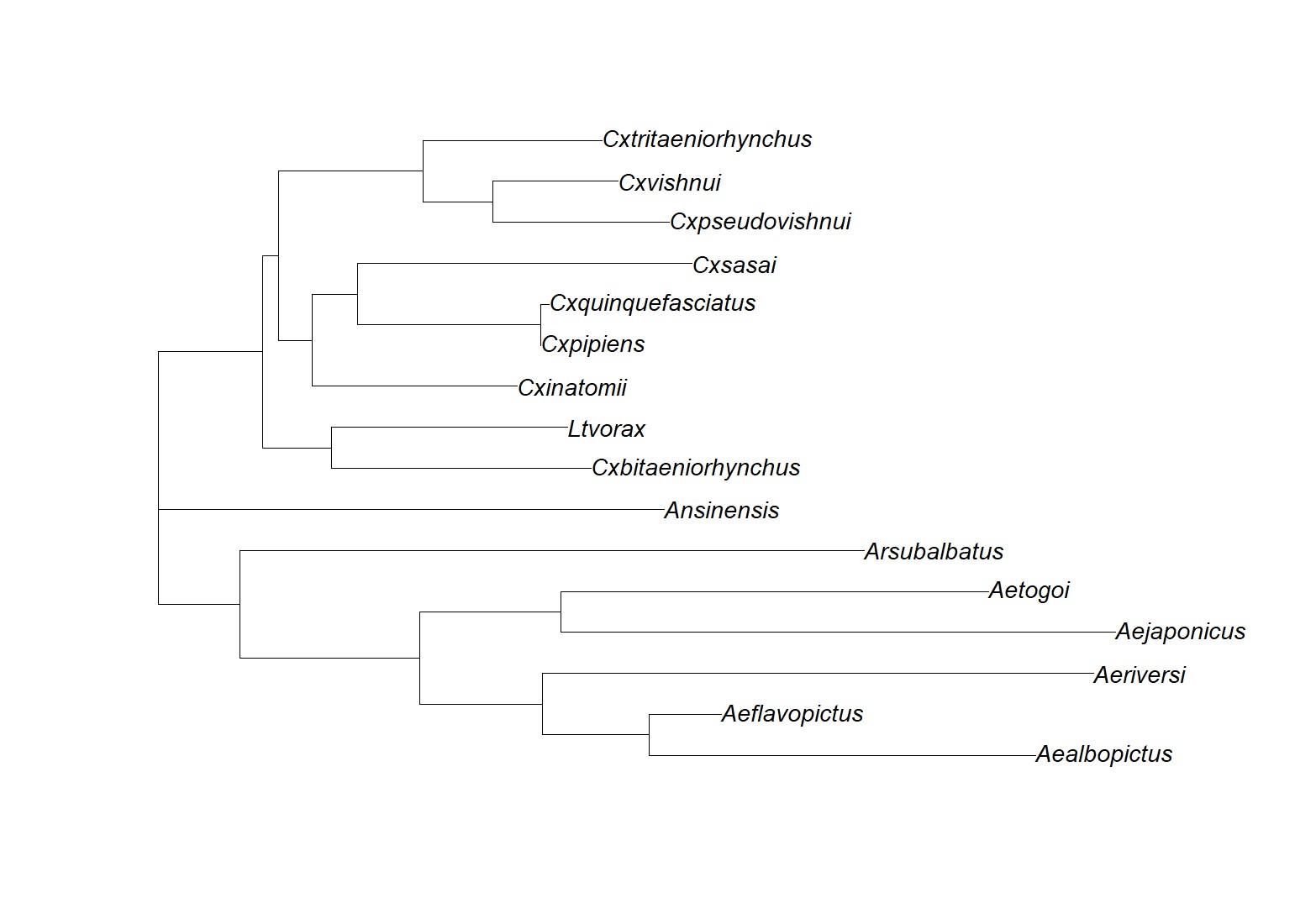

### Supplementary Fig. 2

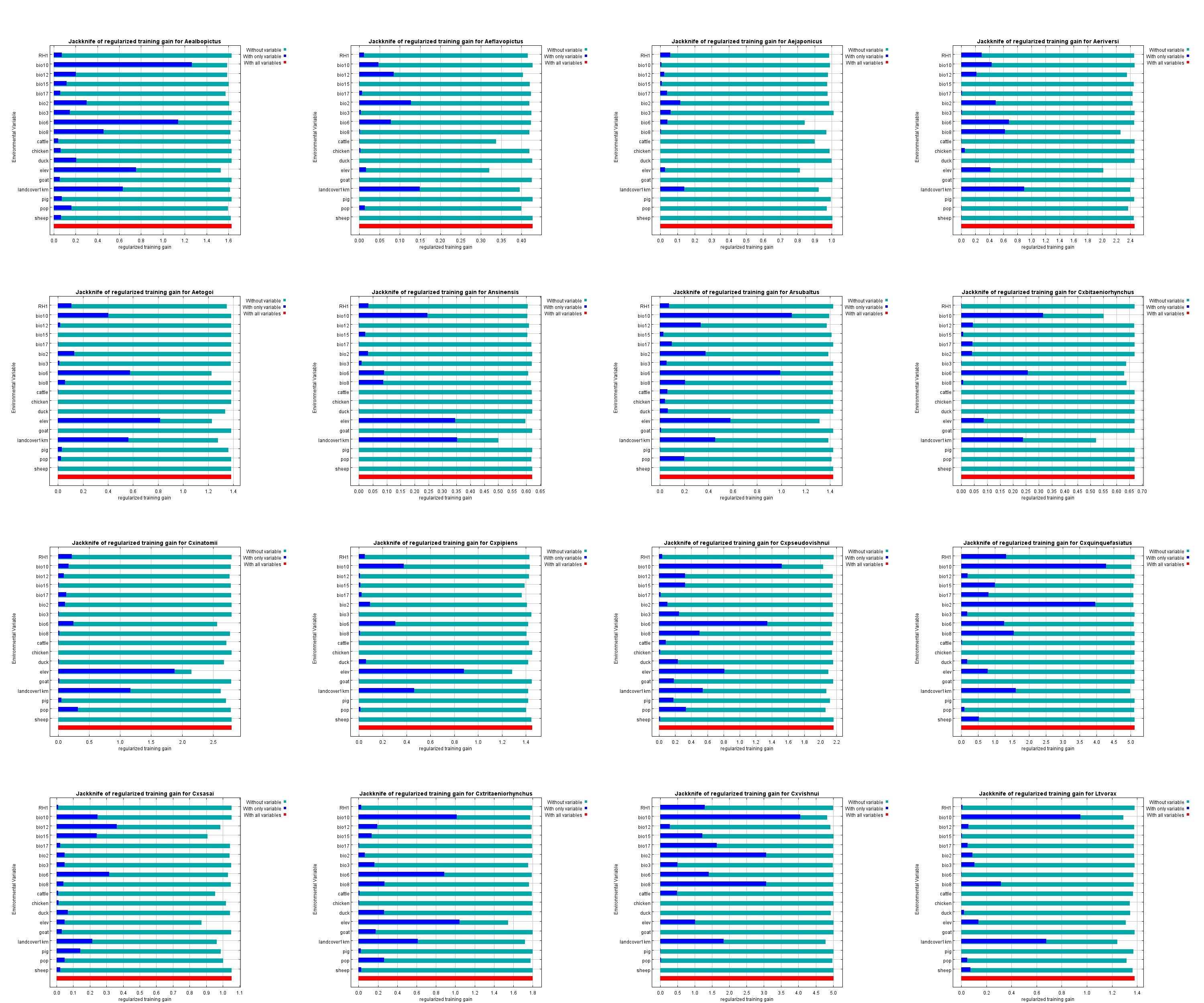

### Supplementary Fig. 3

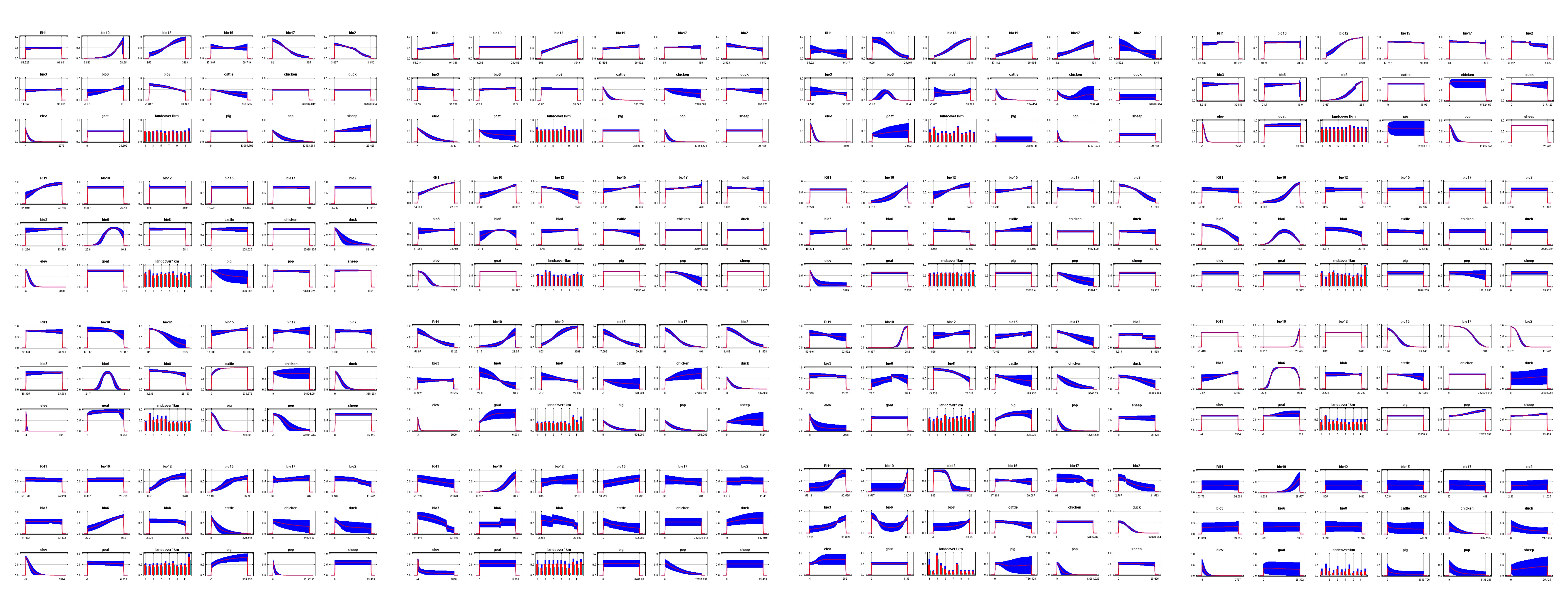

### Supplementary Fig. 4

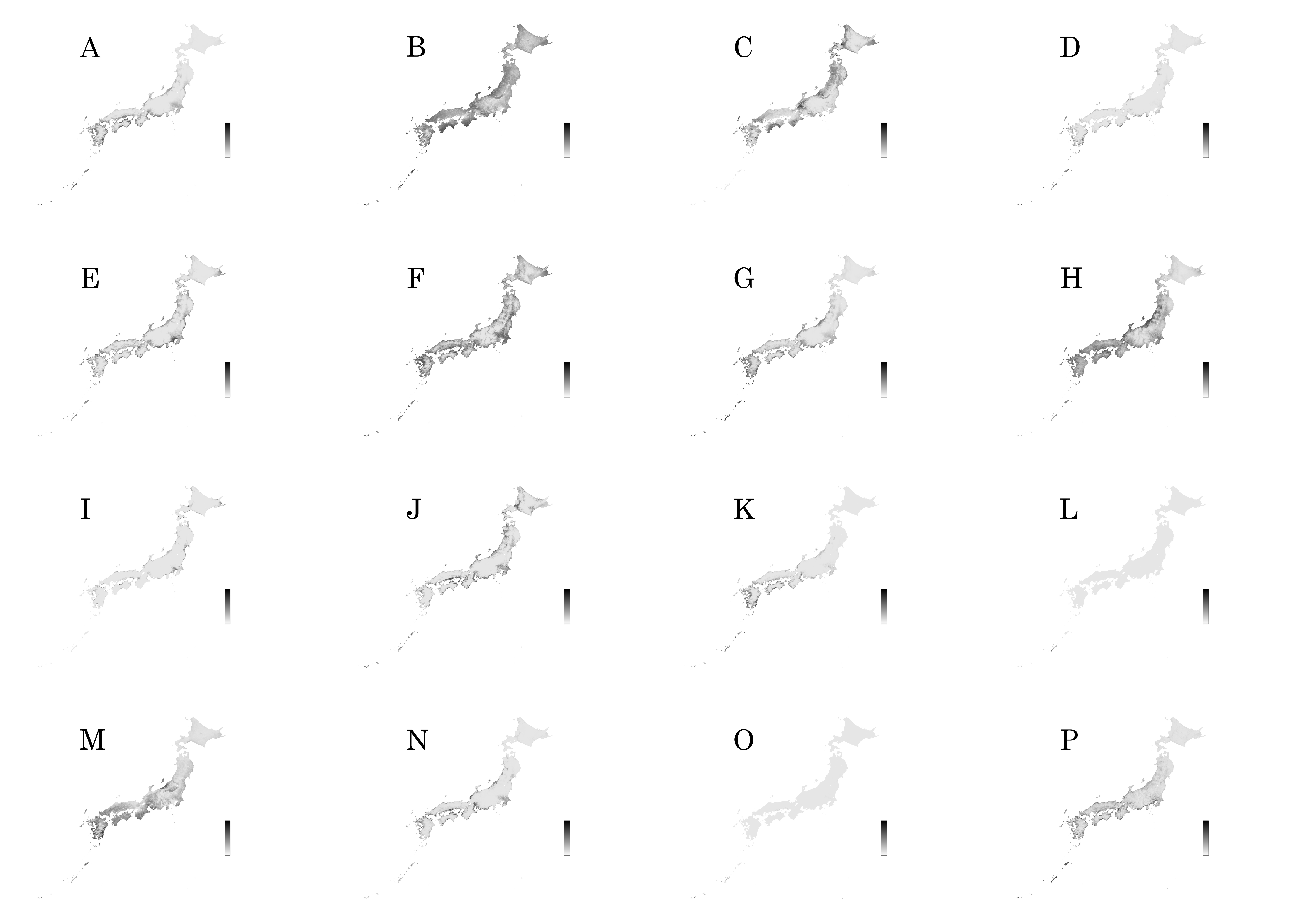

### Supplementary Fig. 5

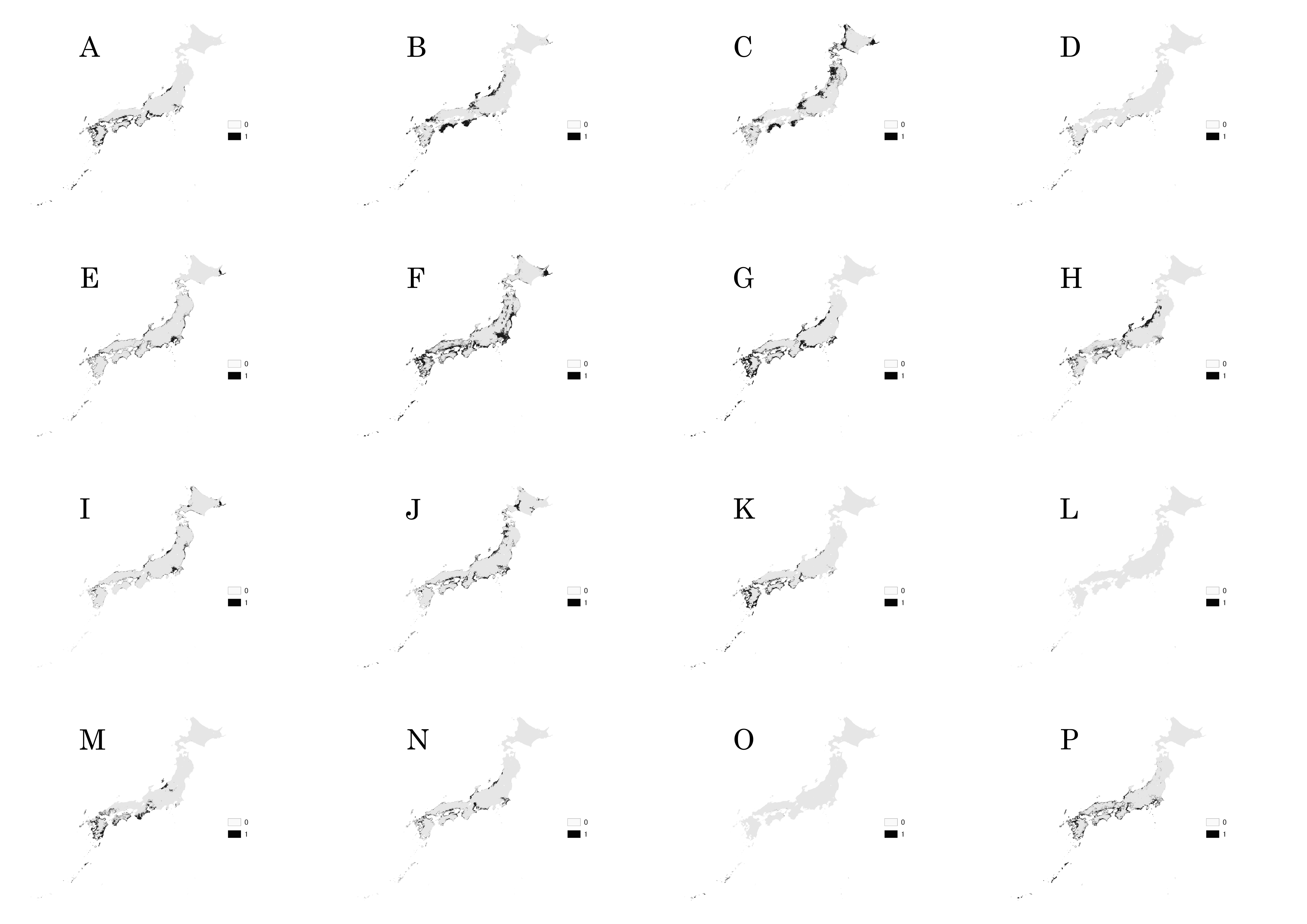
